# Soil protist diversity in the Swiss western Alps is better predicted by topo-climatic than by edaphic variables

**DOI:** 10.1101/571760

**Authors:** Christophe V.W. Seppey, Olivier Broennimann, Aline Buri, Erika Yashiro, Eric Pinto-Figueroa, David Singer, Quentin Blandenier, Edward A.D. Mitchell, Hélène Niculita Hirzel, Antoine Guisan, Enrique Lara

## Abstract

**Aim:** General trends in spatial patterns of macroscopic organisms diversity can be reasonably well predicted from correlative models, using for instance topo-climatic variables for plants and animals allowing inference over large scales. By contrast, soil microorganisms diversity is generally considered as mostly driven by edaphic variables and, therefore, difficult to extrapolate on a large spatial scale based on predictive models. Here, we compared the power of topo-climatic vs. edaphic variables for predicting the diversity of various soil protist groups at the regional scale.

**Location:** Swiss western Alps.

**Taxa:** Full protist community and nine clades belonging to three functional groups: parasites (Apicomplexa, Oomycota, Phytomyxea), phagotrophs (Sarcomonadea, Tubulinea, Spirotrichea) and phototrophs (Chlorophyta, Trebouxiophyceae, Bacillariophyta).

**Methods:** We extracted soil environmental DNA from 178 sites along a wide range of elevations with a random-stratified sampling design. We defined protist Operational Taxonomic Units assemblages by metabarcoding of the V4 region of the ribosomal RNA small sub-unit gene. We assessed and modelled the diversity (Shannon index) patterns of all selected groups as a function of topo-climatic and edaphic variables using Generalized Additive Models.

**Results:** The respective significance of topo-climatic and edaphic variables varied among taxonomic and – to a certain extent – functional groups: while many variables explained significantly the diversity of phototrophs this was less the case for parasites. Generally, topo-climatic variables had a better predictive power than edaphic variables, yet predictive power varied among taxonomic and functional groups.

**Main conclusions:** Topo-climatic variables are, on average, better predictors of protist diversity at the landscape scale than edaphic variables, which opens the way to wide-scale sampling designs avoiding costly and time-consuming laboratory protocols. However, predictors of diversity differ considerably among taxonomic and functional groups; such relationships may be due to direct and/or indirect, e.g. biotic influences. Future prospects include using such spatial models to predict hotspots of diversity or pathogens outbreaks.

## Introduction

Protists, i.e. all eukaryotes with the exception of fungi, plants and animals are hyper-diverse in soil systems (Geisen et al., 2018; Mahé et al., 2017), where they play many ecological roles as primary producers, saprotrophs, predators, or parasites (Adl & Gupta, 2006; Geisen et al., 2016). Photosynthetic groups are essential components of cryptogamic crusts (Elbert et al., 2012; Pushkareva, Johansen, & Elste, 2016) and constitute a significant source of organic carbon for soil organisms (Schmidt, Dyckmans, & Schrader, 2016; Seppey et al., 2017). Predatory protists occupy different levels of the microbial food web, as primary consumers of algae (cyanobacteria or eukaryotic), fungi and bacteria (Bonkowski & Clarholm, 2012; Dumack, Mueller, & Bonkowski, 2016; Hess & Melkonian, 2014), but also occupy higher trophic levels by predating on phagotrophic protists or even micro-Metazoa (e.g. nematodes) (Geisen et al., 2015; Gilbert, Amblard, Bourdier, Francez, & Mitchell, 2000). Parasites are thought to regulate natural populations, notably of animals (Mahé et al., 2017) and can be either very specific such as between the parasitic Gregarines and their animal hosts (Clopton, 2009), or generalist as for Phytomyxea species which can infect hosts from different eukaryotic kingdoms (Neuhauser, Kirchmair, Bulman, & Bass, 2014). Characterizing such complex communities is essential to understand the main on-going ecological processes in soil. This task has been rendered possible only recently with the development of high throughput sequencing, allowing to assess the taxonomic diversity of soil protists, to infer functional diversity and to determine how the patterns and drivers of this diversity compare to the better-known plants and animals.

As a whole, soil protist communities have been shown to respond to edaphic condition such as gradients of pH (Dupont, Griffiths, Bell, & Bass, 2016), nutrients and moisture (Singer et al., 2018) as well as pesticide amounts (Ekelund, 1999; Foissner, 1999; Nesbitt & Adl, 2014) and other perturbations (Foissner, 1997). These variables are rarely integrated in spatial modelling of biodiversity in general Mod, Scherrer, Luoto, & Guisan, (2016) for plant communities), especially at broad spatial scales, because they are most often not available at the sites of species observations and not easily generalizable in a spatially-explicit way (Buri et al., 2017; Dubuis et al., 2013). On the other hand, topo-climatic variables (such as slope steepness or air temperature) can be more easily modelled at large spatial scales using digital elevation models based on interpolations of weather stations and/or remote sensing methods. These variables already proved useful to model the spatial distribution of plants and animals (Franklin, 2010; Guisan, Thuiller, & Zimmermann, 2017; Peterson et al., 2011) but, to our knowledge, very rarely on micro-organisms. As a consequence, spatial modelling of the distribution of microorganisms has been restricted to small areas or aquatic environments (Bulit, 2014; Fraile, Schulz, Mulitza, & Kucera, 2008; King et al., 2010; Langer, Weinmann, Loetters, Bernhard, & Roedder, 2013; Mitchell et al., 2000; Zaric, Schulz, & Mulitza, 2006; Zinger, Shahnavaz, Baptist, Geremia, & Choler, 2009). The development of such models at the landscape scale would allow assessing at a much broader scale the processes driven by micro-organisms, such as nutrient cycling, predicting zones at risk of pathogenic outbreaks or simply identifying protist diversity hotspots.

Here, we built spatial predictive models of protist diversity, focussing on general communities as well as on nine broad protist taxa chosen within three functional groups - phototrophs, phagotrophs and parasites - along a wide elevation gradient in the western Swiss Alps. We assessed the diversity of protists in 178 meadow soil samples, resulting from a robust random-stratified field survey by metabarcoding of the V4 regions of the small sub-unit rRNA gene. This study assessed the extent of protist diversity in mountainous meadows and determined to what extent two sets of environmental variables (edaphic and topo-climatic) can predict this diversity over the whole Swiss western Alps of the Vaud state. In addition, we brought an interpretation of the patterns observed based on knowledge of the lifestyles of the different groups surveyed.

## Material and Methods

### Sampling

Meadow soils were sampled from 194 plots distributed across the Swiss western Alps; of these plots, 178 samples successfully yielded sequencing data and were used in the current study. Sampling was performed from July 4^th^ to September 1^st^ 2013 according to a random stratified sampling design. From each plot, five soil cores (100 grams per core between the depths of 0-5 cm after removing plants, mosses, and insects) were taken from the four corners and the centre of a 2 m^2^ plot. The five cores, were then pooled in a sterile plastic bag and kept in an icebox or at 4 °C until DNA extraction and soil analyses were done. A subsample of the pooled soil was also flash frozen at each sampling site and kept frozen until further soil analyses. For more details, see Yashiro et al. (2016).

### Edaphic variables

We selected eight edaphic variables that were measured directly in the field or on the soil samples. Soil temperature was measured in the field. The relative humidity (rh) was assessed by weighing the mass of the soil sample before and after drying at 105 °C during 2 days. Soil organic Carbon content was determined by loss of ignition (LOI) at 1050 °C. The percentage of shale was determined by laser granulometry. The pH and electro-conductivity (EC) were measured from a soil and Milli-Q water slurry in a 1:2.5 and 1:5 (wt/vol) ratio respectively. Total phosphorus amount was determined by colorimetric analysis after a mineralisation at 550 °C with Mg(NO_3_)_2_. The C/N ratio was calculated from the total organic carbon and nitrogen percentages measured by ROCK EVAL pyrolysis (Vinci Technologies, Ruell-Malmaison, France) and combustion infrared spectroscopy (Carlo Erba CNS2500 CHN), respectively. See Yashiro et al. (2016) and Buri et al. (2017) for more details.

### Topo-climatic variables

Values for seven topo-climatic variables were retrieved from maps of 25 square meter resolution for each sample location. We used the number of growing degree-days above 0 °C (gdd), potential evapo-transpiration (etp), topography (topo), slope southness (asp) and slope steepness (slp)(Zimmermann & Kienast, 1999; Zimmermann, Edwards, Moisen, Frescino, & Blackard, 2007). In addition, we calculated the summer temperature average (tmean678) and precipitation sum (psum678) for the months of June to August with values of monthly temperature means and precipitation sums from 1981 to 2010. See Buri et al. (2017) for more details.

### Molecular analysis

DNA was extracted from the soil samples using the MoBio PowerSoil DNA extraction kit (Calsbad, CA, USA) following the manufacturer instructions. The V4 region of the 18S rRNA gene was then amplified using the general eukaryotic primers TAReuk454FWD1 and TAReukREV3 (CCAGCASCYGCGGTAATTCC / TYRATCAAGAACGAAAGT) (Stoeck et al., 2010). The PCR mix was composed of 3 µL DNA extract, 0.4 µL of 10 mg/mL BSA, 4 µL of PCR buffer (Promega GoTaq M7845), 0.2 µL of Taq polymerase (Promega GoTaq M7845), 0.6 µL of dNTPs (Promega kit U1420), 0.6 µL of each primer (MicroSynth, Balgach, Switzerland), and 10.6 µL of ultra-pure water. The PCR reactions started with a denaturation step at 95 °C for 5 min followed by 45 cycles of 94 °C for 30 s, 47 °C for 45 s and 72 °C for 1 min, and terminated with an elongation step of 72 °C for 10 min. For each DNA sample, the amplifications were performed in triplicate with a PTC-200 Peltier Thermo Cycler (BioConcept, Allschwil, Switzerland). DNA was then quantified with a Qubit^®^ 2.0 Fluorometer (Invitrogen) and 20 ng of each triplicate were pooled. A DNA library was prepared from the pools using the TruSeq Nano PCR-free Library Preparation kit and the paired-end 2×300 bp sequencing was done on an Illumina^®^ MiSeq at the University of Geneva (Molecular Systematics & Environmental Genomics Laboratory).

### Bioinformatics pipeline

Good quality sequences were selected based on their nucleotides phred scores. Every sequence with a phred score average below 20 for a 50 nucleotides window was discarded. The chimeras were then removed using the program vsearch 1.11.1 (Rognes, Flouri, Nichols, Quince, & Mahé, 2016) by comparing the environmental sequences 1) with each other for each replicate and 2) against the PR^2^ database trimmed according to the V4 primers (downloaded on the 12 September 2016; Guillou et al. (2013)). To reduce the noise caused by very rare sequences, we then discarded every singleton. Triplicates were then pooled according to their respective samples and OTUs were built with the program swarm 2.1.8 (Mahé, Rognes, Quince, de Vargas, & Dunthorn, 2015). The dominant sequence of each OTU was taxonomically assigned by aligning it to the trimmed PR^2^ database using the global pairwise alignment program ggsearch 36.3.6 (Pearson, 2000).

We removed every OTU that did not belong to protists, namely Metazoa, Embryophyceae and Fungi. We also discarded OTUs with a percentage of identity (PID) below 65% with the database PR^2^ as sequences with such low PID are usually of prokaryotic origin (threshold verified manually by aligning low PID environmental sequences on GenBank database). From the 178 plots, 4 were sampled twice and 13 were sampled three times during the sampling period. For each of these 17 plots we took the average (2 samples) or median (3 samples) sequence abundance of each OTU for the samples from the same plot. In addition of the total protist community matrix, we also selected nine broad taxonomic groups (I.e. clades, low taxonomic resolution) from three functional groups (1) parasites: Apicomplexa, Oomycota, Phytomyxea; (2) phagotrophs: Sarcomonadea, Tubulinea, Spirotrichea and (3) phototrophs: Chlorophyceae, Trebouxiophyceae, Bacillariophyta). These taxa were selected because they are abundant and diverse in soils and are functionally homogeneous. For each of these taxa, we established a PID threshold verified manually on GenBank to discarded potential misidentification (Apicomplexa: 80%, Oomycota: 80%, Phytomyxea: 75%, Sarcomonadea: 80%, Tubulinea: 75%, Spirotrichea: 90%, Chlorophyceae: 90%, Trebouxiophyceae: 85%, Bacillariophyta: 77%).

### Richness and diversity analyses

For each of the ten taxonomic data sets (all protists plus nine broad groups), OTU richness and Shannon diversity (H) were calculated, and the differences between their statistical distributions tested by a multiple comparisons of mean rank sums test (Nemenyi test; Hollander, Wolfe, & Chicken, 2015, `posthoc.kruskal.nemenyi.test` function, ‘PMCMR’ package 4.1; Pohlert, 2014). Computation of H indices includes quantitative data, classically proportion of a given species in a given sample, which can be reasonably inferred by numbers of reads in High Throughput Sequencing data. Indeed, there is a correspondence between this number of reads and the biovolume of individual organisms that has been showed for many groups of protists (Giner et al., 2016; Kosakyan, Mulot, Mitchell, & Lara, 2015; Weber & Pawlowski, 2013). H indices provide thus a reasonable estimation of the OTU diversity. We then assessed the relationships between Shannon diversity (H), and topo-climatic and edaphic variables. For this, we firstly assessed pairwise correlations between all predictors and, in all pairs with correlation >0.7, we only kept the expectedly most causal one for further analyses to avoid collinearity issues (Dormann et al., 2013) (see Appendix Fig. S1.1 in Supporting Information). Then, for each of the ten data sets, H was modelled as a function of the environmental variables using a Generalized Additive Model (GAM; assuming Gaussian residuals and identity link function). For each data set, three models were calibrated; the first with topo-climatic variables only, the second with edaphic variables only, and the third with both sets of variables. All models were iterated 100 times based on bootstraps composed of 80% of the 178 original samples. In total 10×3×100 models were fitted. For each model, the predictive power was estimated as the Root Mean Square Error (RMSE) calculated on the independent samples not included to build the model (20% left-out samples). The effect of taxonomic group and the set of predictors on predictive power (RMSE) was tested by a Nemenyi test. Finally, the diversities of the nine broad taxa and total protist diversity were extrapolated on the full area of the western Swiss Alps with a GAM including the topo-climatic variables (i.e. the only spatially-explicit variables).

## Results

### Observed diversity patterns

We retrieved a total of 24’322’487 good quality sequences of which 97% were not chimeric and 71% were not singletons. The 17’110’114 remaining sequences were clustered into 41’048 OTUs of which 19’260 were assigned to protists (Table 1). Protist diversity was dominated (proportion of sequences) by Cercozoa, (principally Sarcomonadea and Thecofilosea), and Alveolata of which more than half were assigned to Apicomplexa and ca. 45% to Ciliophora (mostly from classes Spirotrichea, Oligohymenophorea, Litostomatea and Colpodea) (see Appendix Fig. S1.2). The three other dominant groups were the Stramenopiles (including Oomycota and Bacillaryophyta), Amoebozoa (including Tubulinea) and Archaeplastida (with Chlorophyceae and Trebouxiophyceae) (see Appendix Fig. S1.2).

**Table 1:**
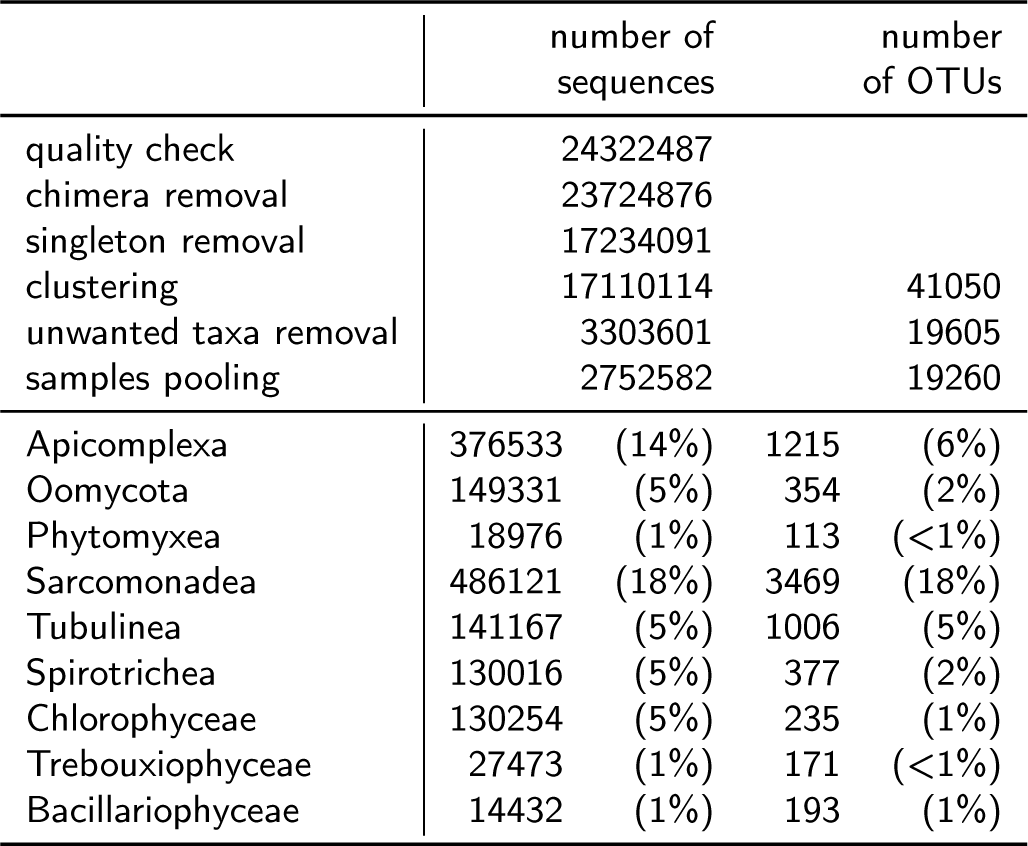
Number of sequences and operational taxonomic units retrieved from 178 plots in the Swiss western Alps, through each step of the analysis for the total community and for the nine broad taxa. The numbers between parenthesis represent the percentage of the total community.

The nine chosen taxa jointly contributed to over half (54%) of all retained sequences and represented over 35% of the total OTU richness (Table 1). The average richness per sample of these clades varied from 7 (Phytomyxea) to 249 (Sarcomonadea). Richness was in average lowest for phototrophs (15 OTUs / sample) and highest for phagotrophs (122 OTUs / sample; Fig. 1). Shannon diversity indices followed the same trend, varying from an average value of 1.1 (Phytomyxea) to 4.3 (Sarcomonadea).

**Figure 1:**
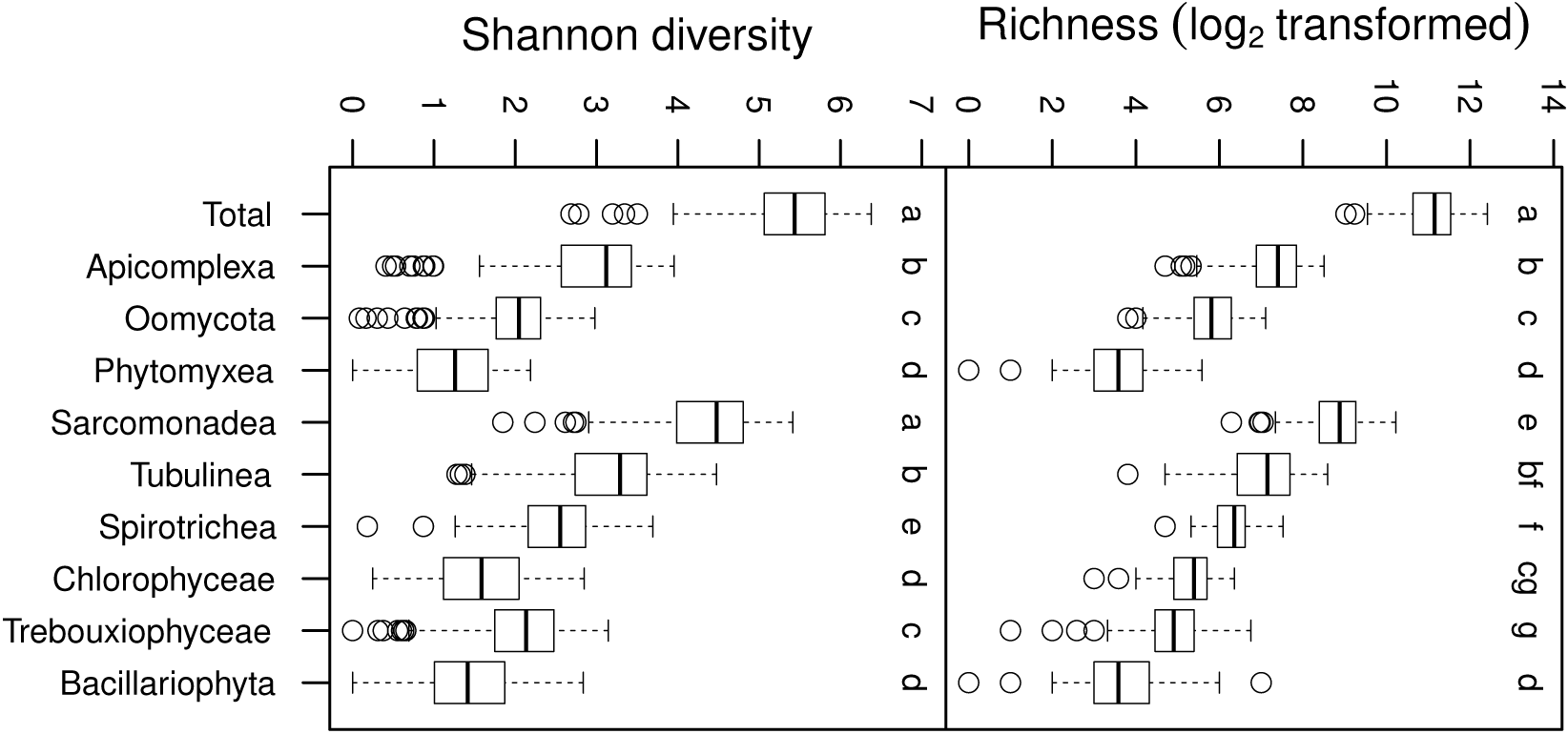
Shannon diversity and richness (log transformed) distributions of protist operational taxonomic units communities retrieved from 178 plots in the Swiss western Alps. The distributions are shown for the total community as well as for nine broad taxa. The letters above the boxplots represent groups according to a multiple comparison mean rank sums test (Nemenyi test P < 0.05).

### Environmental models of diversity

The percentage of variance (adjusted-R^2^) of the Shannon diversity in the total and broad taxonomic groups explained by the combination of both topo-climatic and edaphic variables ranged from 6% (Bacillariophyta) to 33% (Chlorophyceae) (Table 2). The environmental variables explaining a significant (p<0.05) portion of the protist diversity in these models with combined datasets were: slope steepness (in 4 taxa), pH (3 taxa), mean summer temperature (2 taxa), Soil Organic C (2 taxa), shale percentage (1 taxon), C/N (1 taxon), phosphorus (1 taxon) and EC (1 taxon) (Table 2). The predictive power showed lower RMSE values (i.e. a better power) for the topo-climatic than for the edaphic variables for all taxa except the Chlorophyceae, Trebouxiophyceae and Sarcomonadea where the values were higher or similar (Fig. 2). In addition, the RMSE of the models calculated on the edaphic and topo-climatic variables together were never significantly lower than the RMSE calculated for the topo-climatic variable alone. The RMSE also varied among taxonomic groups and the diversity of certain taxa were significantly better predicted (e.g. Oomycota) than others (e.g. Apicomplexa) (Table 3). This RMSE variation was also observed at the functional level (see Appendix Table S1.1): the RMSE of functional group were always as good as or better than the RMSE calculated from the total community.

**Table 2:**
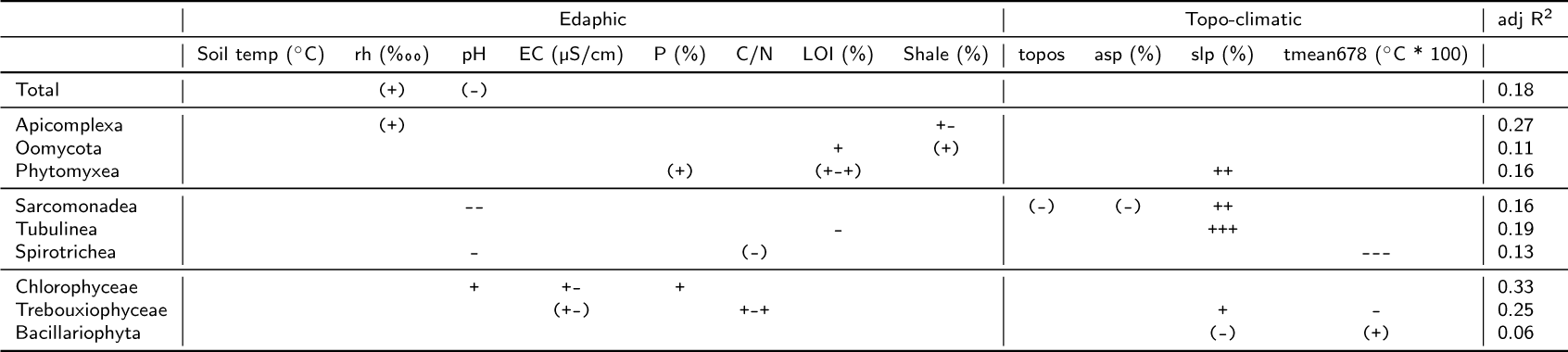
Significance of edaphic (soil temperature: Soil temp, relative humidity: rh, pH, electroconductivity: EC, total phosphorus amount: P, carbon/nitrogen ratio: C/N, loss of ignition: LOI, shale percentage) and topo-climatic (topography: topo, slope southness: asp, slope steepness: slp, summer temperature average: tmean678) predictors on the diversity modelled (Generalized Additive Model) from total micro-eukaryotic community and nine broad taxonomic groups from operational taxonomic units gathered from 178 meadow soils in the Swiss western Alps. The + and – signs show if the diversity is positively or negatively associated to the predictor and the number of signs inform on the strength of the association (between parenthesis: P < 0.1, one sign: P < 0.05, two signs: P < 0.01, three signs: P < 0.001). The -+ and +-indicate minimum and maximum of diversity at mid-predictor value respectively. Details of the response of each taxonomic group to the different variables can be found in Fig. S1.3.

**Table 3:** Average and standard deviation of the predictive power (Root Mean Square Error: RMSE) of edaphic, topo-climatic and overall predictors calculated on the diversity of protist operational taxonomic units from the total community and nine broad taxa as well as the functional groups the taxa were belonging to. The communities were retrieved from 178 meadow soils in the Swiss western Alps. The RMSE were calculated on 100 cross validation of Generalized Additive Models performed with 20% of the samples as test dataset. The letters between parenthesis represent significantly different groups according to a multiple comparison mean rank sums test (Nemenyi test P < 0.05) for each of the total communities and nine broad taxa or for the total community and the three functional groups.

|  | edaphic |  | topo-climatic |  | overall |  |
| --- | --- | --- | --- | --- | --- | --- |
| Total | 0.69 ± 0.10 | (a) | 0.61 ± 0.11 | (ab) | 0.66 ± 0.11 | (ab) |
| Apicomplexa | 0.73 ± 0.11 | (a) | 0.67 ± 0.12 | (c) | 0.71 ± 0.11 | (ac) |
| Oomycota | 0.53 ± 0.07 | (b) | 0.43 ± 0.06 | (d) | 0.45 ± 0.06 | (d) |
| Phytomyxea | 0.62 ± 0.07 | (c) | 0.58 ± 0.06 | (ab) | 0.62 ± 0.07 | (be) |
| Sarcomonadea | 0.62 ± 0.09 | (c) | 0.59 ± 0.10 | (ab) | 0.60 ± 0.09 | (e) |
| Tubulinea | 0.69 ± 0.07 | (a) | 0.57 ± 0.06 | (ae) | 0.61 ± 0.07 | (be) |
| Spirotrichea | 0.63 ± 0.08 | (c) | 0.54 ± 0.06 | (e) | 0.60 ± 0.07 | (e) |
| Chlorophyceae | 0.53 ± 0.06 | (b) | 0.57 ± 0.06 | (ae) | 0.55 ± 0.06 | (f) |
| Trebouxiophyceae | 0.61 ± 0.10 | (c) | 0.60 ± 0.06 | (ab) | 0.62 ± 0.10 | (be) |
| Bacillariophyta | 0.70 ± 0.07 | (a) | 0.62 ± 0.07 | (bc) | 0.70 ± 0.08 | (c) |
| Total | 0.69 ± 0.10 | (a) | 0.61 ± 0.10 | (a) | 0.66 ± 0.10 | (a) |
| Parasites | 0.63 ± 0.10 | (b) | 0.56 ± 0.10 | (b) | 0.59 ± 0.10 | (b) |
| Phagotrophs | 0.65 ± 0.09 | (c) | 0.57 ± 0.08 | (b) | 0.60 ± 0.08 | (bc) |
| Phototrophs | 0.61 ± 0.10 | (b) | 0.59 ± 0.07 | (a) | 0.62 ± 0.10 | (c) |

**Figure 2:**
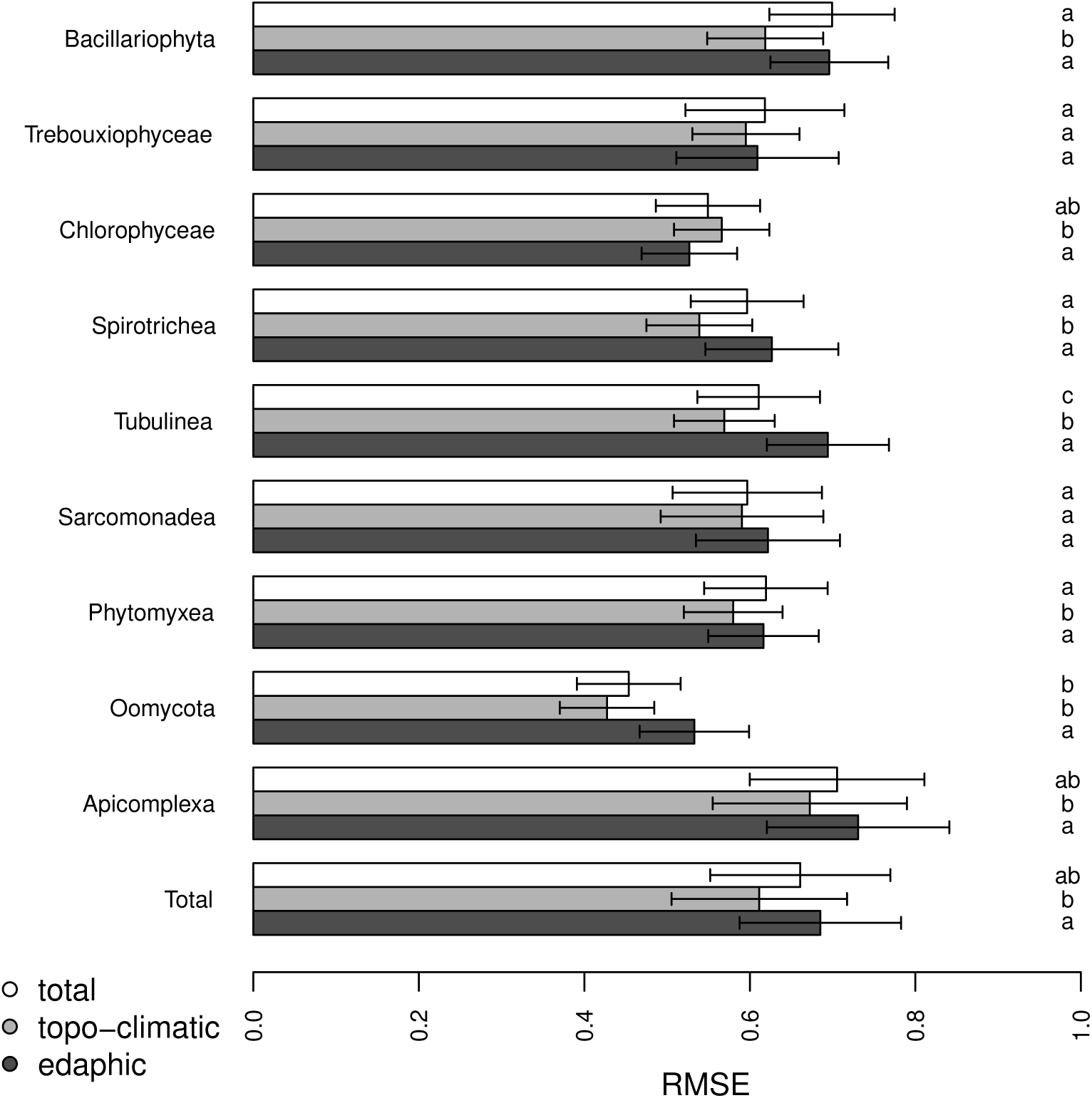
Predictive power (Root Mean Square Error: RMSE) of edaphic (dark grey), topo-climatic (pale gray) and overall (white) predictors calculated on the diversity of protist operational taxonomic units from the total community and nine broad taxa retrieved from 178 meadow soils in the Swiss western Alps. The RMSE were calculated on 100 cross validation of Generalized Additive Models performed with 20% of the samples as test dataset. The letters on the right of the barplot represent significantly different groups according to a multiple comparison mean rank sums test (Nemenyi test P < 0.05) for each of the total communities and nine broad taxa.

## Discussion

### General patterns of micro-eukaryotic diversity in soils

Our study revealed several important findings on patterns of protist diversity across temperate mountain landscapes. Phagotrophs (e.g. Sarcomonadea & Tubulinea) and parasites (Apicomplexa) were the most abundant functional groups in terms of read abundance. Apicomplexan sequences, albeit numerous, were much less abundant and diversified than in Neotropical soils: as arthropods are less abundant and diversified in temperate regions, this brings further support to the hypothesis that soil apicomplexan diversity mirrors that of arthropods in the ecosystem (Mahé et al., 2017). Another abundant parasitic group is the Oomycetes (Stramenopiles), which contains many plant parasites, but also animal pathogens and a few free-living, saprotrophic forms (Beakes, Glockling, & Sekimoto, 2012; Lara & Belbahri, 2011). Oomycetes were shown to be common and diverse in temperate soil systems (Seppey et al., 2017; Singer et al., 2016). By contrast they are less abundant and diverse in neotropical forest soil ecosystems, where they comprise mostly animal parasites (Mahé et al., 2017).

Within phagotrophs, the high proportion of sequences from Cercozoa (mostly to Sarcomonadea) was in line with previous soil eukaryotic DNA surveys (Bates et al., 2013; Harder et al., 2016; Seppey et al., 2017). Earlier studies based on microscopical observations showed the prevalence of these groups in soils (Adl & Gupta, 2006). Ciliates were also a well-represented phagotrophic group, and were dominated by Spirotrichea, which corroborates also other findings on soil protist molecular diversity (Lara, Berney, Ekelund, Harms, & Chatzinotas, 2007). In summary, the protist communities found in the Swiss western Alps are typical for average soil ecosystems and the findings can probably be extrapolated to other regions.

### Model fit and predictive power of topo-climatic and edaphic variables

Slope steepness and pH were the two variables most often found to significantly contribute to the fit of our different protist diversity models. Slope steepness affects drainage and leaching of nutrients and is generally inversely correlated to soils depth. Nevertheless, a enhanced drainage reduces the likelihood of water-logging which would select for very specialized protists tolerating anoxia and generally would lead to lower diversity. Soil pH is well known as a major driver of microbial diversity, including bacteria (Santoyo, Hernandez-Pacheco, Hernandez-Salmeron, & Hernandez-Leon, 2017; Yashiro et al., 2016), fungi (Noyce et al., 2016; Pellissier et al., 2014; Zhang, Jia, & Yu, 2016) and protists (Bates et al., 2013; Dupont et al., 2016). The relationship between pH and protist diversity was significant only for three groups, being negative for two groups of phagotrophs (Spirotrichea and Sarcomonadida) and positive for Chlorophyceae. It is unclear if these relationships reflect a direct effect of pH or rather indirect effects such as biotic effects (e.g. impact on bacterial or fungal food sources), the availability of nutrients for the growth of autotrophs, or other drivers.

Predictability varies also to a large extent between functional groups. Indeed, while many variables explained significantly the diversity of phototrophs and phagotrophs, it was less so for parasites (see Appendix Table S1.1). The latter depend only indirectly on environmental values, and mainly on their hosts, which brings logically supplementary noise in analyses. For nine out of the ten taxonomic group tested, the predictive power of the topo-climatic variables was either significantly better, or at least not different than the ones including the edaphic variables. Moreover, it was never lower than the predictive power of the models including both sets of variables. This suggests that, within the levels of predictability achieved, predictive models built solely on topo-climatic variables are as accurate, or possibly even better than the models built with the addition of edaphic variables. These variables are available at large scales and are already largely used for modelling the spatial distribution of macroorganisms (Guisan & Zimmermann, 2000), to the contrary of local edaphic values that are always tedious and costly to measure in the landscape across large regions and environmental gradients. These findings open the way to larger sampling designs that could further increase the performance of models.

The correspondence between OTUs and biological species has always been a hot topic in eukaryotic environmental microbiology. Indeed, a single SSU rRNA gene sequence may include, in certain groups, a wide diversity of species with different lifestyles and ecological preferences. This has been shown for different soil protists such as ciliates (Lara, & Acosta-Mercado, 2012); in contrast, in Myxomycetes (Amoebozoa), SSU sequences are truly hypervariable and discriminate relatively accurately between species (Dahl et al., 2018). There is, therefore, no general rule that applies to all eukaryotes. However, SSU sequences are generally considered good proxy for eukaryotic diversity, as they were the first benchmark for protist barcoding (Pawlowski et al., 2012), and therefore, predictions based this proxy can as well be considered a good estimation of actual protists’ diversity.

### Interpretation of the spatial patterns of protist diversity modelled with topo-climatic variables

As for macro-organisms (D’Amen, Pradervand, & Guisan, 2015; Dubuis et al., 2011; McCain, 2005; Reymond, Purcell, Cherix, Guisan, & Pellissier, 2013), but unlike other soil micro-organisms (Bryant et al., 2008; Fierer et al., 2010; Pellissier et al., 2014), protists diversity show clear spatial and elevational patterns when only topo-climatic variables are taken to build the model (Fig. 3). This patterns seems to be driven by summer temperature in most cases (see Appendix Table S1.1), either in a positive (Bacillariophyta, Phytomyxea and Tubulinea), unimodal (Apicomplexa, Sarcomonadea and Spirotrichea) or negative way (Chlorophyceae, Oomycota). A positive correlation of diversity with temperature (and, thus, productivity) is a typical pattern in macroecology that can be related to the species-energy hypothesis as long as moisture is not a limiting factor (Fernández et al., 2016), or other models for diversity patterns (Huston, 1994; see Spehn & Körner, (2009) for elevation gradients). On the other hand, if moisture is limiting, unimodal patterns are to be expected, and diversity peaks where both moisture and energy are optimal (water energy model: Fernández et al. (2016)) intermediate disturbance hypothesis or mid-domain effect (discussed for the same area in Dubuis et al. (2011)). Finally, Chlorophyceae and Oomycota are typically sensitive to high temperatures and desiccation, both including often flagellated life stages for dispersal that needs at least a thin water film to disperse (Jeger & Pautasso, 2008). In addition, Chlorophyta high diversity in the lowest temperature zone (Fig. 3) could be explained by the fact that micro-eukaryotic algae have a higher growth rate at low temperatures, favouring diversification in cold environments (Rose & Caron, 2007) or possibly reduced competition from vascular plants.

**Figure 3:**
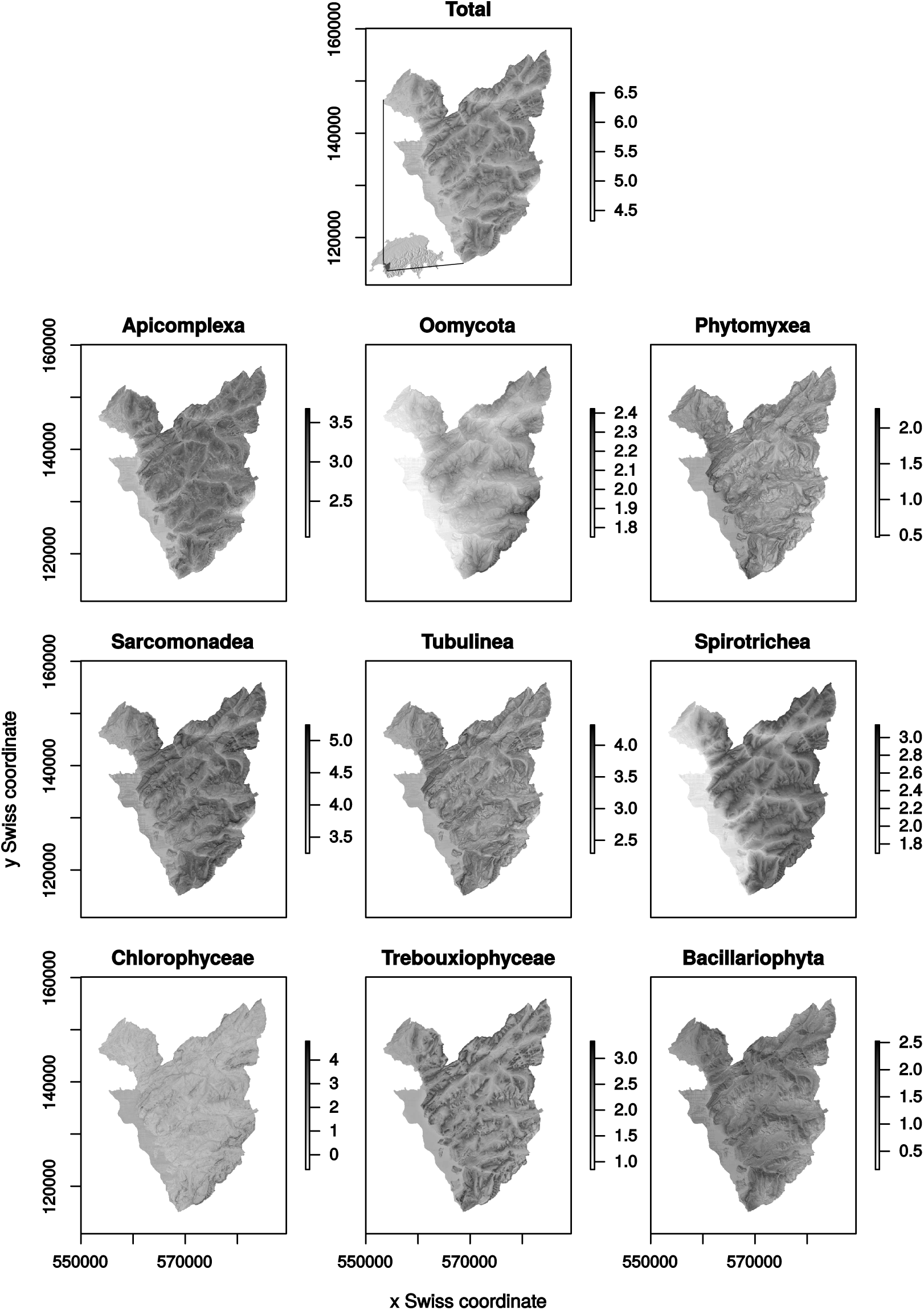
Diversity of the total protist community and nine broad taxa predicted from Generalized Additive Model through the Swiss western Alps based on the topography, slope southness, slope steepness and average temperature from June to August.

## Conclusion

We showed that the diversity of some taxa and functional groups, is explained up to >30% by topo-climatic and edaphic conditions. A somewhat surprising result is that topography and climate predicted protist diversity as well or better than the edaphic variables more commonly used in soil microbial studies. This implies that soil protist diversity patterns could be at least partly inferred, for some groups (e.g. Chlorophyceae) and to some extent (22%), based on topo-climatic spatial models only.

Such an approach could be applied at finer taxonomic levels to predict the distribution of individual species, which would be of high socio-economic relevance in the case of invasive agricultural or forestry pests of economic importance such as certain oomycetes. The models could be improved by refining the taxonomic groups, as taxa responding more homogeneously to environmental conditions may show stronger correlation with abiotic variables than the broad group classification we used. For instance, the Oomycota contain organisms belonging to other functional groups than parasites (e.g. saprotroph; Beakes et al., 2012; Lara & Belbahri, 2011) or able to target a wide range of hosts (e.g. *Phytophothora cinnamomi*; Hardham, 2005). These improvements would pave the way toward extrapolation of protists diversity across large spatial scales and provide useful tools to identify biodiversity hotspots, predict spatially the risk of pathogen infection or model soil protist diversity according to future environmental change scenarios.

## Acknowledgements

The authors would like to thank all fields and laboratory assistants and technicians who participate in the alpine soil project, with a particular thank to Amandine Pillonel, Laura Desponds and Dessislava Savova Bianchi for the laboratory work. Special thanks to the Transports Publics du Chablais and Glacier 3000 (https://www.glacier3000.ch/en) who provide their installations for free to carry the soils from the sampling sites. We would also like to thanks all anonymous farmers who allowed sampling on their lands.

The study was founded by the Swiss National Science Foundation under the projects 310003A 143960 to E.L. and 31003A-152866 (SESAM’ALP), PDFMP3-135129 (MICROBIAL BIOGEOGRAPHY) and CR23I2-162754 (INTEGRALP) to A.G., as well as the internal funding of the Universities of Neuchâtel and Lausanne. E.L. would also like to thanks the program “Atracción de talentos” from the Community of Madrid project 2017-T1/AMB-5210. E.Y. also thanks the European Community FP7-PEOPLE-2010-IIF program (MP-Alps, grant agreement 273965), the Agassiz Foundation, and the Pro-Femmes Fellowship program from the Faculty of Biology and Medicine of the University of Lausanne.

A significant part of the computations was also performed on resources provided by the Calculations Center of the Faculty of Science of the University of Neuchâtel and by UNINETT Sigma2 - the National Infrastructure for High Performance Computing and Data Storage in Norway.

## Biosketch

The Laboratory of Soil Biodiversity (https://www.unine.ch/biolsol), led by Prof. Edward A.D. Mitchell, is interested in the diversity, biogeography and ecology of soil organisms with a strong focus on protists and links to other soil organisms and ecosystem ecology. The lab combines observational and experimental studies leading to applications in biomonitoring, palaeoecology, ecotoxicology and forensic sciences. The Spatial Ecology Group (http://www.unil.ch/ecospat), led by Prof. Antoine Guisan, is specialized in spatial modeling of biodiversity at the levels of species, communities, and ecosystems. Models are applied to the conservation of endangered species, the management of biological invasions, and the assessment of global change impact on biodiversity, with a special and long-term focus on above-and below-ground biota in the Western Swiss Alps.

## Author contributions

E.Y., E.A.D.M., H.N.H., A.G., and E.L. conceived the idea; E.Y., E.A.D.M., A.G. and E.L. provided the funding; A.B., E.Y., E.P.F. and A.G. collected the data; A.B., E.Y., E.P.F., D.S., Q.B. and E.L. performed the laboratory work; C.V.W.S., O.B., A.B., A.G. and E.L. analysed the data and C.V.W.S., O.B., E.Y., D.S., Q.B., E.A.D.M., A.G. and E.L. wrote the manuscript. Al authors gave final approval for publication.

