## Supporting Information for "Soil protist diversity in the Swiss western Alps is better predicted by topo-climatic than by edaphic variables"

[1]Laboratory of Soil Biodiversity, University of Neuchâtel, Émile-Argand 11, 2000 Neuchâtel, Switzerland
[2]Department of Arctic and Marine Biology, University of Tromsø, Framstredet 39, 9019 Tromsø, Norway
[3]Department of Ecology and Evolution, University of Lausanne, Biophore, 1015 Lausanne, Switzerland
[4]Institute of Earth Surface Dynamics, University of Lausanne, Géopolis, 1015 Lausanne, Switzerland
[5]Department of Fundamental Microbiology, University of Lausanne, Biophore, 1015 Lausanne, Switzerland
[6]Department of Zoology, Institute of Biosciences, University of São Paulo, Matão 14, 05508-090 São Paulo, Brazil
[7]Real Jardín Botánico, CSIC, Plaza de Murillo 2, 28014 Madrid
[8]Botanical Garden of Neuchâtel, Perthuis-du-Sault 58, 2000 Neuchâtel, Switzerland
[9]Institute for Work and Health, Universities of Lausanne and Geneva, 1066 Epalinges-Lausanne, Switzerland
[+]Co-last authors

SUPPORTING INFORMATION

### APPENDIX S1 Principal Component Analysis calculated on the predictors used to model the diversity of protist taxa through the Swiss western Alps

**Figure S1.1**: Principal Component Analysis calculated on the environmental variables (soil temperature: Soil_temp, electro-conductivity: EC, relative humidity: rh, loss on ignition: LOI, percentage of shale: Shale, pH, phosphorus percentage: P, C/N ratio: C_N, growing degree day: gdd, potential evapotranspiration: etp, topography: topo, slope southness: asp, slope steepness: slp, summer average temperature: tmean678, summer precipitation sum: psum678) measured from 178 plots through the Swiss western Alps. The first and second, and first and third axis are shown. Note the tight correlation between the tmean678 and the variables removed for the final analyses, namely gdd, etp and psum678.

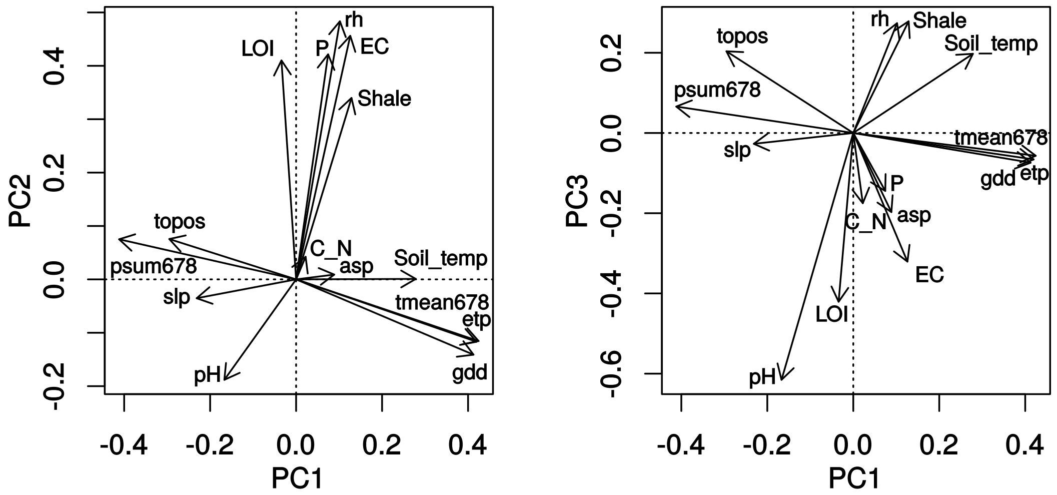

**Figure S1.2**: Four topo-climatic predictors (topography: topo, slope southness: asp, slope steepness: slp, average temperature from June to August from 1981 to 2010: tmean678), spatialized on the area of the Swiss western Alps. These four predictors were used to model the diversity of the total protist community and nine broad taxa through Generalized Additive Models.

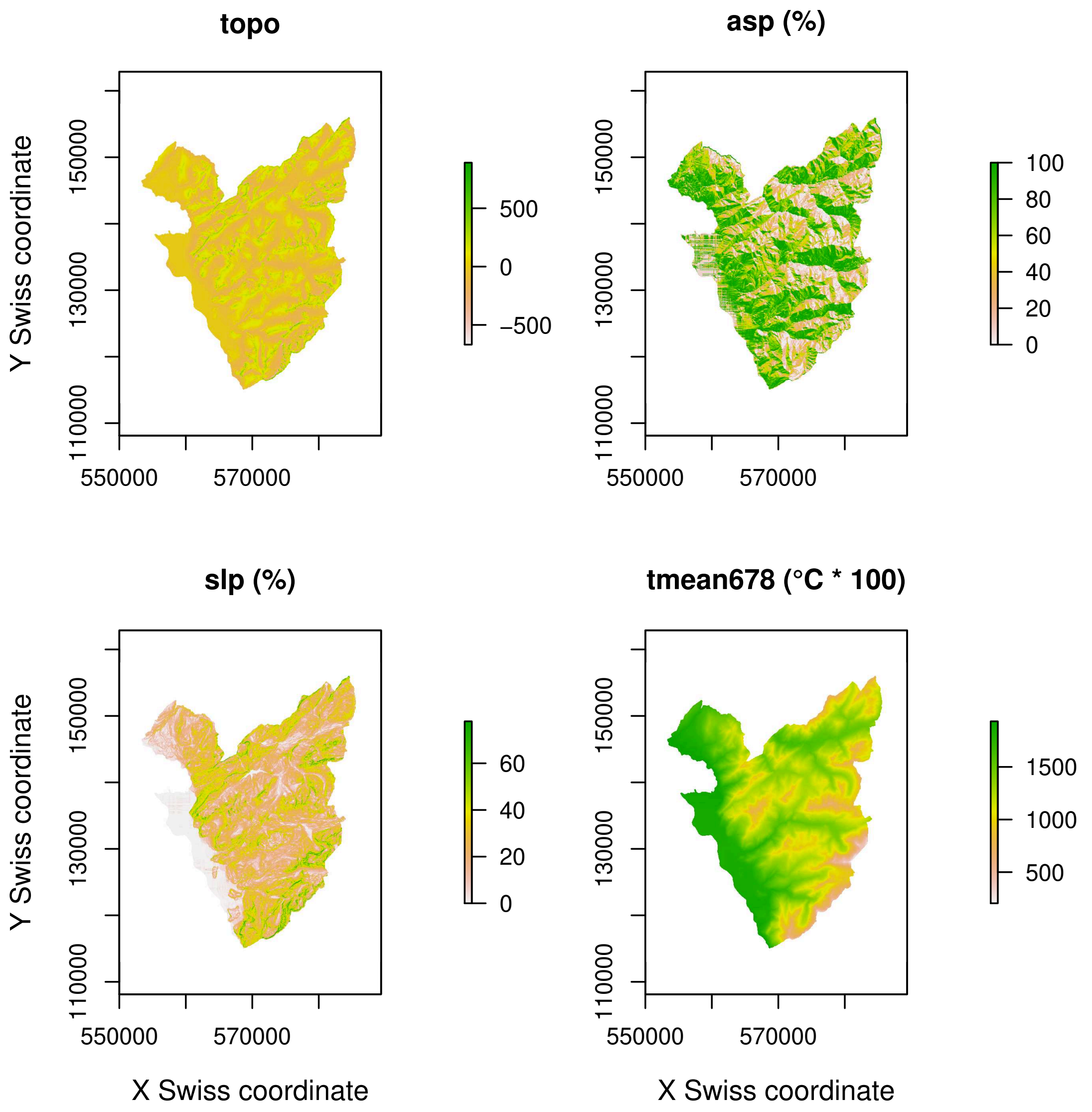

**APPENDIX S2 Relative abundance of protist taxon sequences abundance from 178 soil samples across the Swiss western Alps.**

**Figure S2.1**: Relative abundance of protist taxon sequence abundances from 178 meadow soils from the Swiss western Alps. Only taxa representing at least 1% of the total number of sequences are represented.

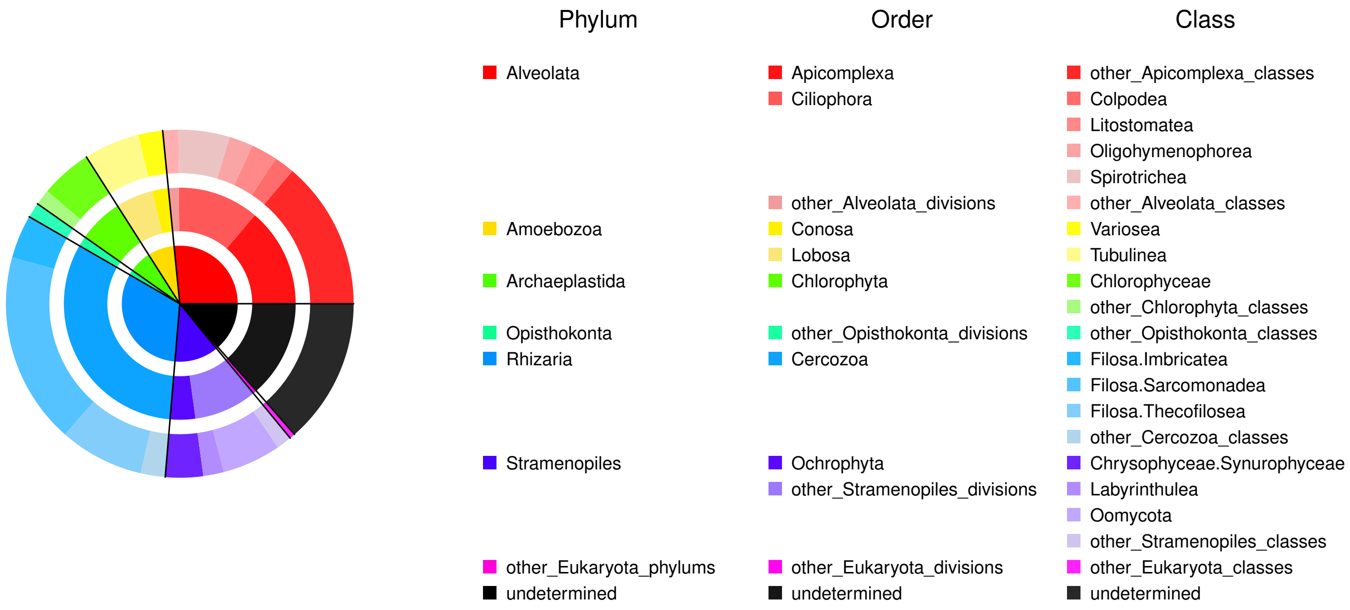

### APPENDIX S3 General Additive Models on protist diversity in function of two sets of predictors (edaphic and topo-climatic)

**Figure S3.1**: Diversity of the total protist community (Total) and all nine broad taxa in function of eight edaphic variables (soil temperature: Soil_temp, residual humidity: rh, pH, electro-conductivity: EC, phosphorus percentage: P, C/N ratio, loss on ignition: LOI and shale percentage: Shale) and four topo-climatic predictors (topography: topo, slope southness: asp, slope steepness: slp, average temperature from June to September: tmean678) through Generalized Additive Models.

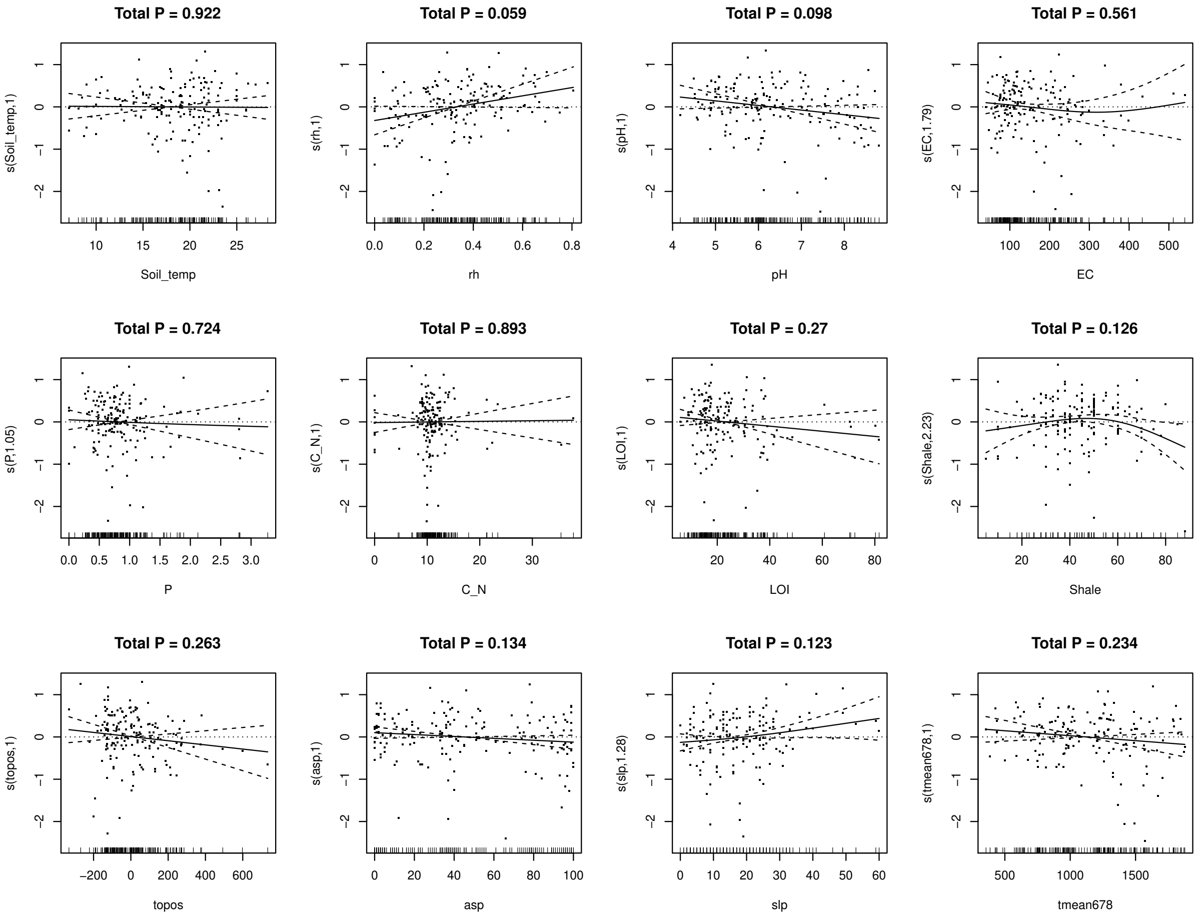

**Figure S3.1**: continuation

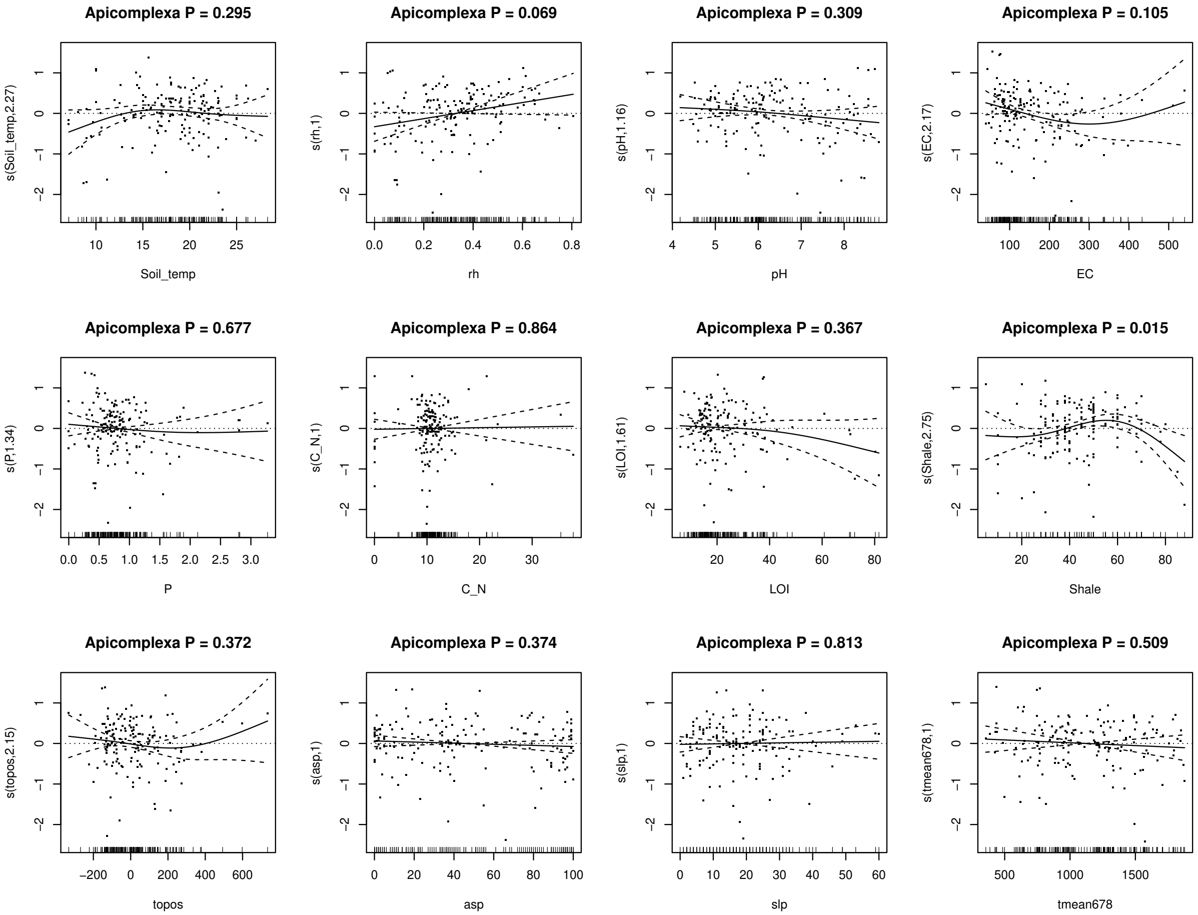

**Figure S3.1**: continuation

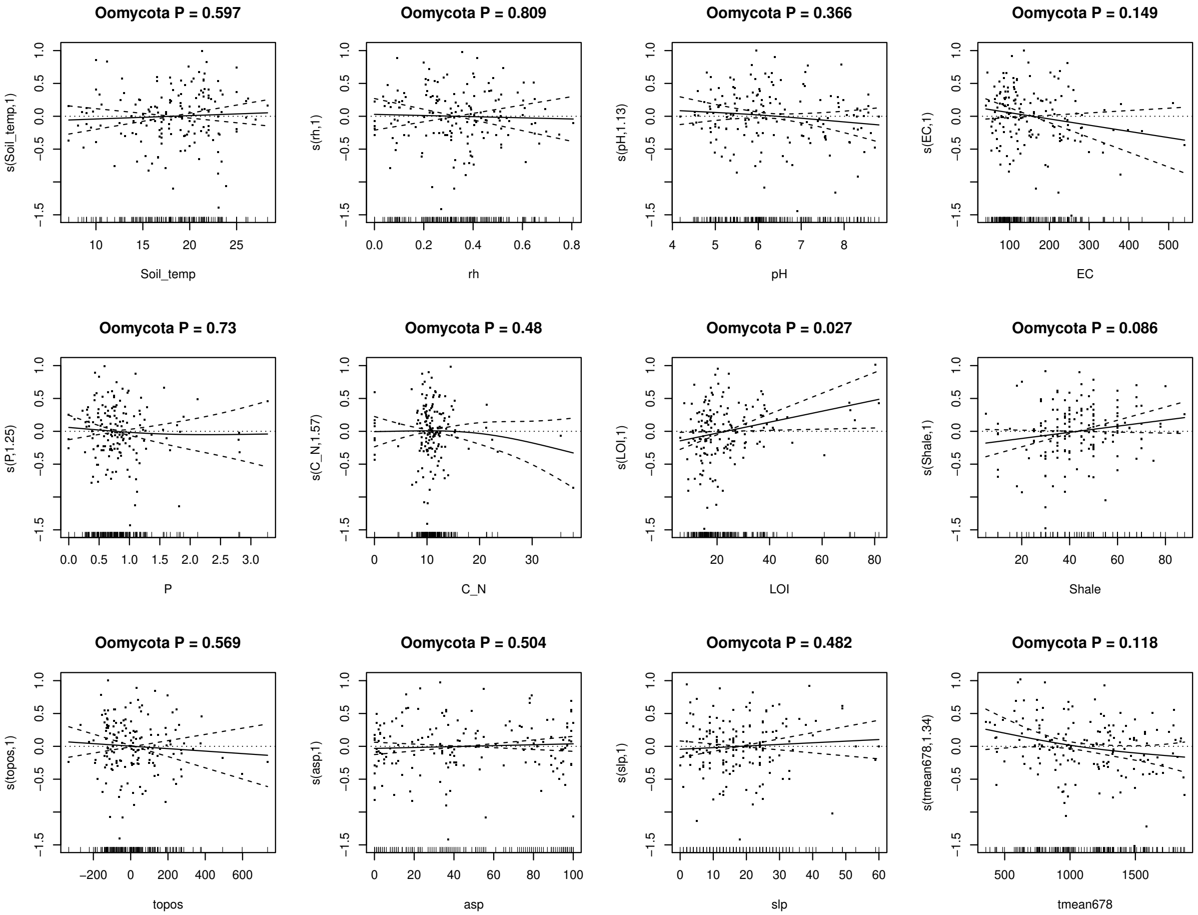

**Figure S3.1**: continuation

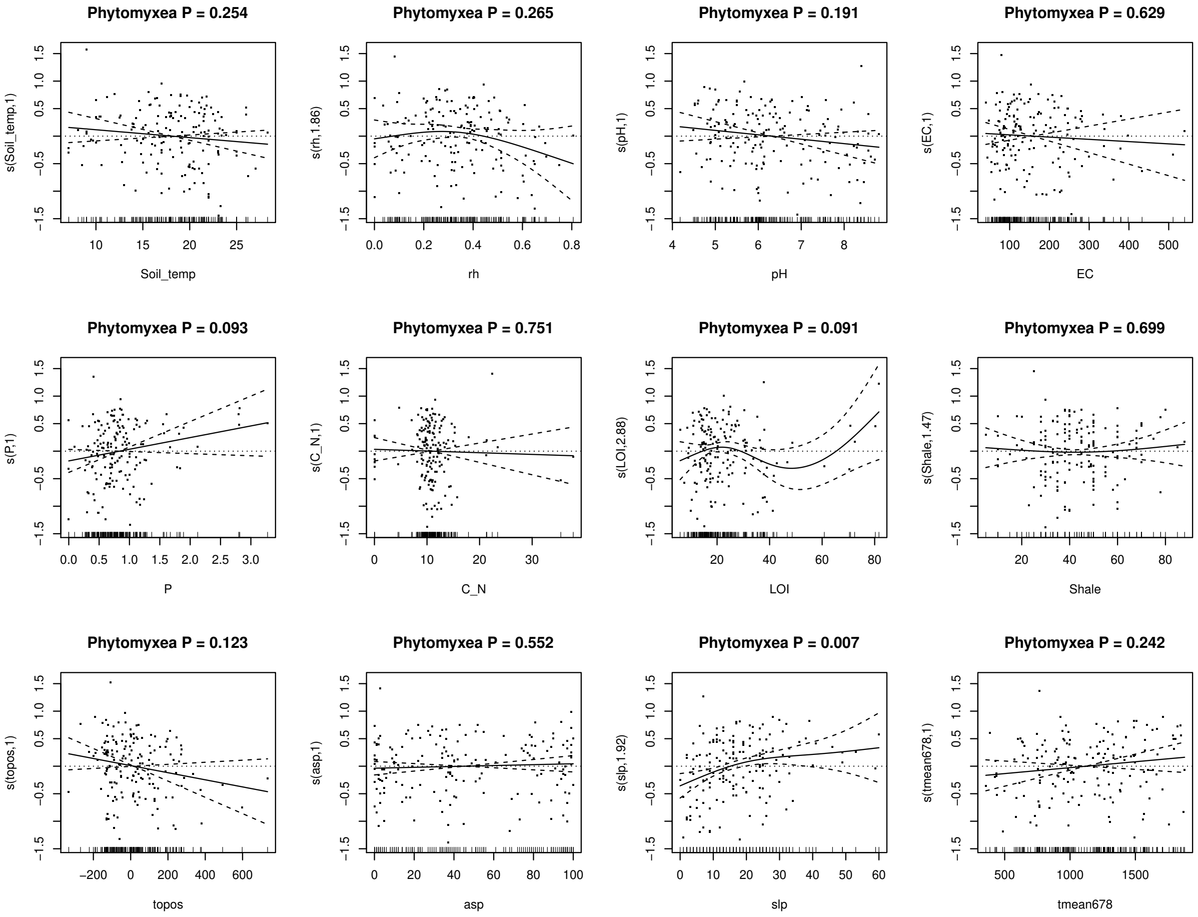

**Figure S3.1**: continuation

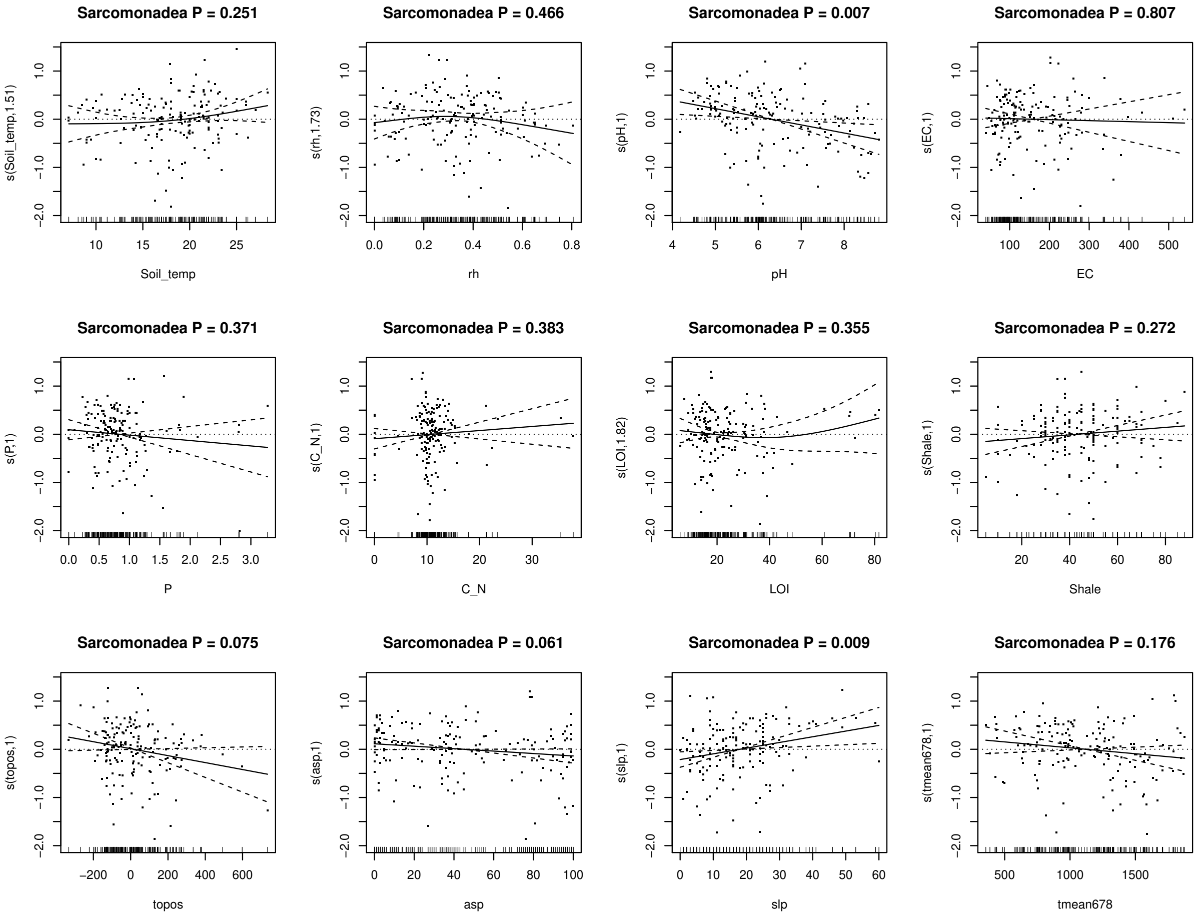

**Figure S3.1**: continuation

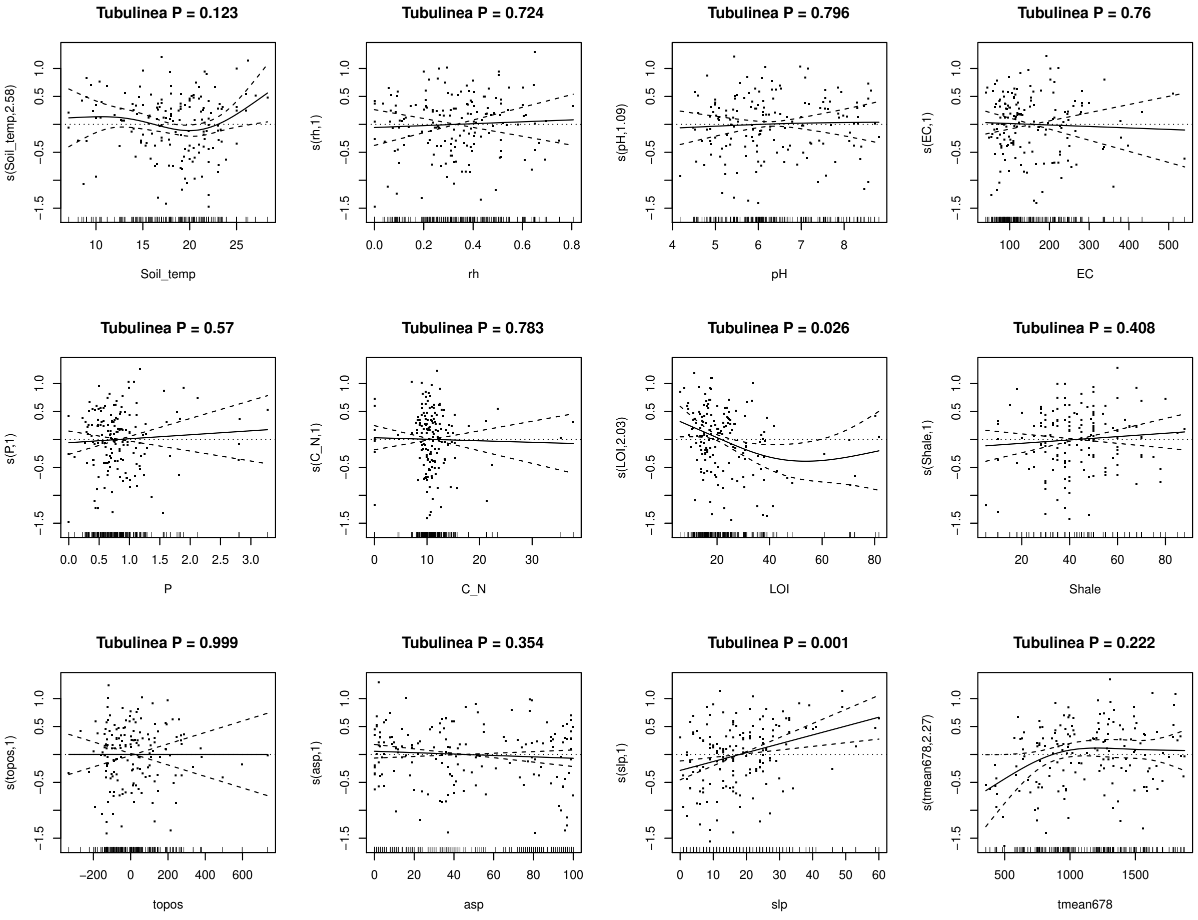

**Figure S3.1**: continuation

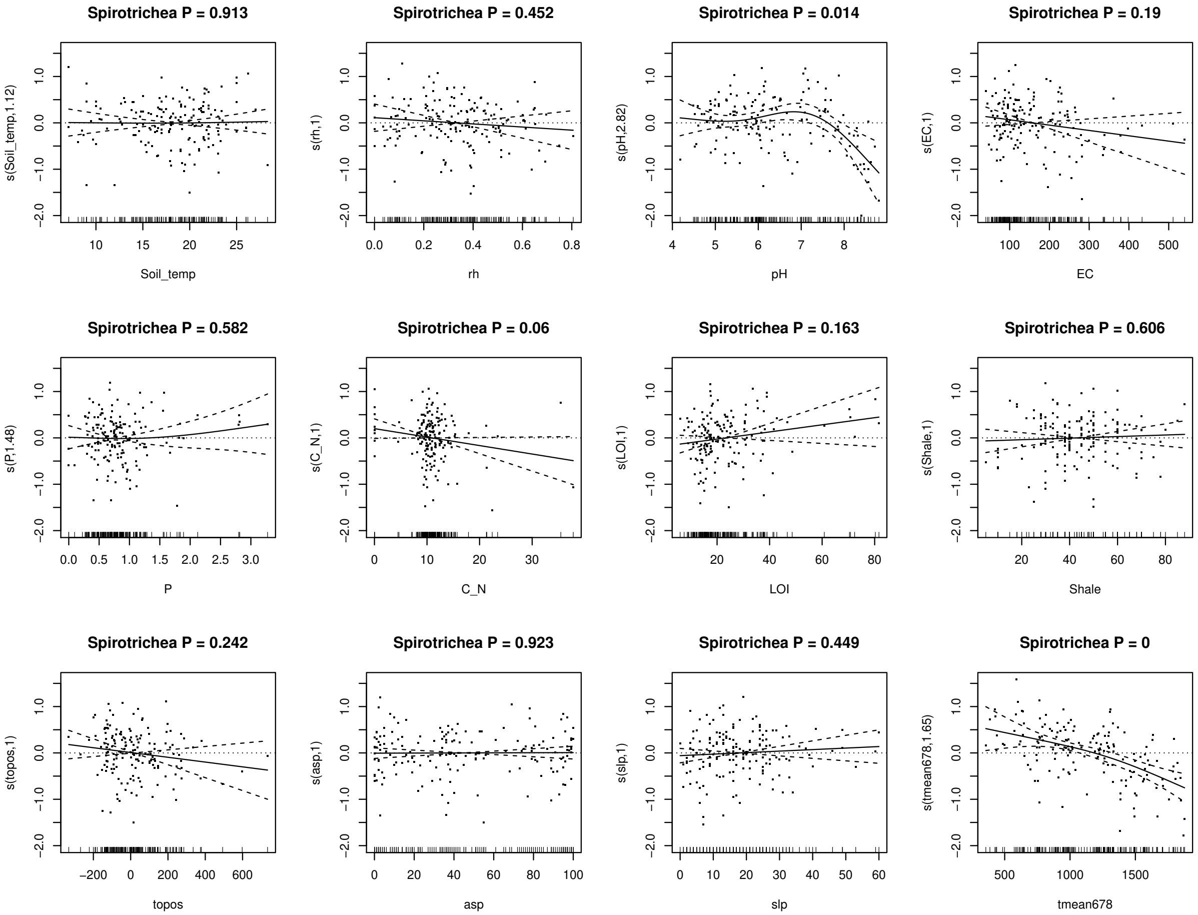

**Figure S3.1**: continuation

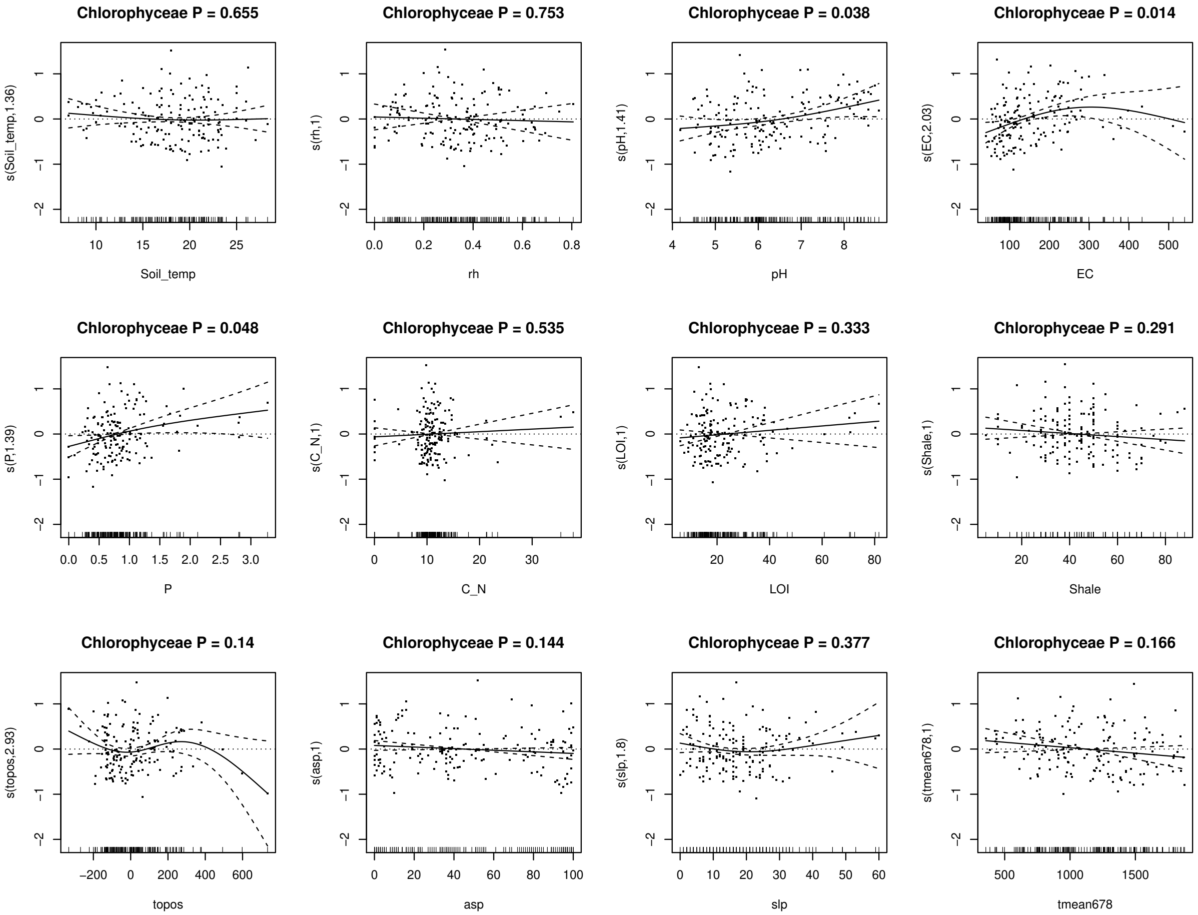

**Figure S3.1**: continuation

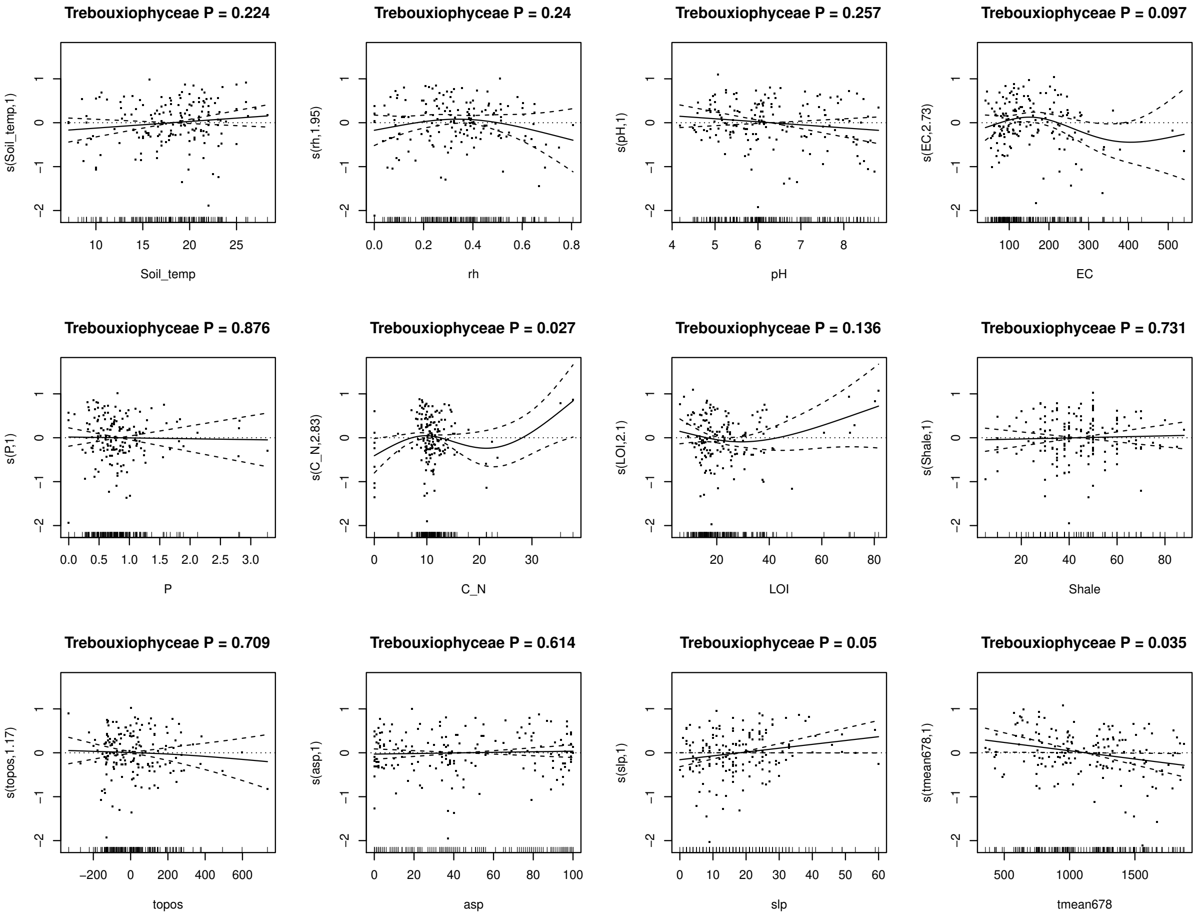

**Figure S3.1**: continuation

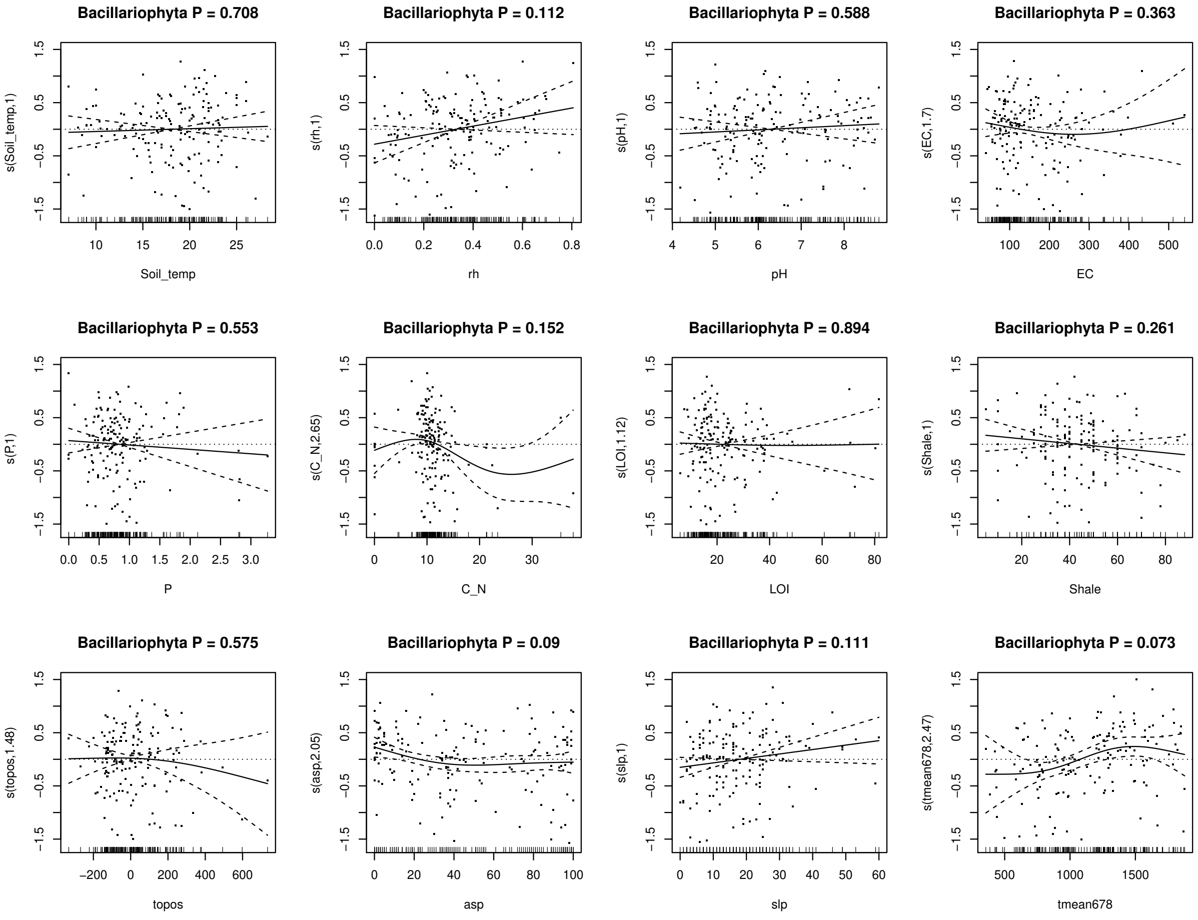

**APPENDIX S4 General Additive Models on protist diversity in function of topo-climatic predictors**

**Table S4.1**: Significance of topo-climatic predictors on the diversity modelled (Generalized Additive Model) from the total micro-eukaryotic community and nine broad taxonomic groups from operational taxonomic units gathered from 178 meadow soils in the Swiss western Alps. The + and - signs indicate if the diversity is positively or negatively associated to the predictor and the number of signs inform on the strength of the association (between parenthesis: P < 0.1, one sign: P < 0.05, two signs: P < 0.01, three signs: P < 0.001). The -+ and +- indicate minimum and maximum of diversity at mid-predictor value respectively. Details of the response of each taxonomic group to the different variables can be found at Figure **S4.2**.

| | Topo-climatic | | | | $R^2$ |
|---|---|---|---|---|---|
| | topos | asp | slp | tmean678 | |
| Total | (+) | (-) | | (++--) | 0.09 |
| Apicomplexa | | | | (+-) | 0.05 |
| Oomycota | | | | - | 0.03 |
| Phytomyxea | | | ++ | + | 0.08 |
| Sarcomonadea | | - | + | +- | 0.08 |
| Tubulinea | | | ++ | + | 0.08 |
| Spirotrichea | | | | +++--- | 0.15 |
| Chlorophyceae | -- | | --++ | -- | 0.22 |
| Trebouxiophyceae | ++-- | | ++ | | 0.11 |
| Bacillariophyta | | (-) | + | ++ | 0.13 |

**Figure S4.2**: Diversity of the total protist community and all nine broad taxa in function of four topo-climatic predictors (topography: topo, slope southness: asp, slope steepness: slp, average temperature from June to September: tmean678) through Generalized Additive Models.

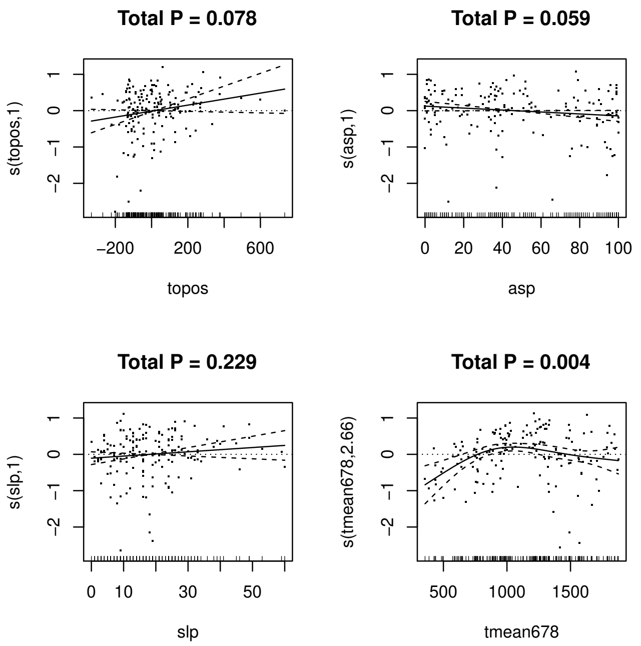

**Figure S4.2**: continuation

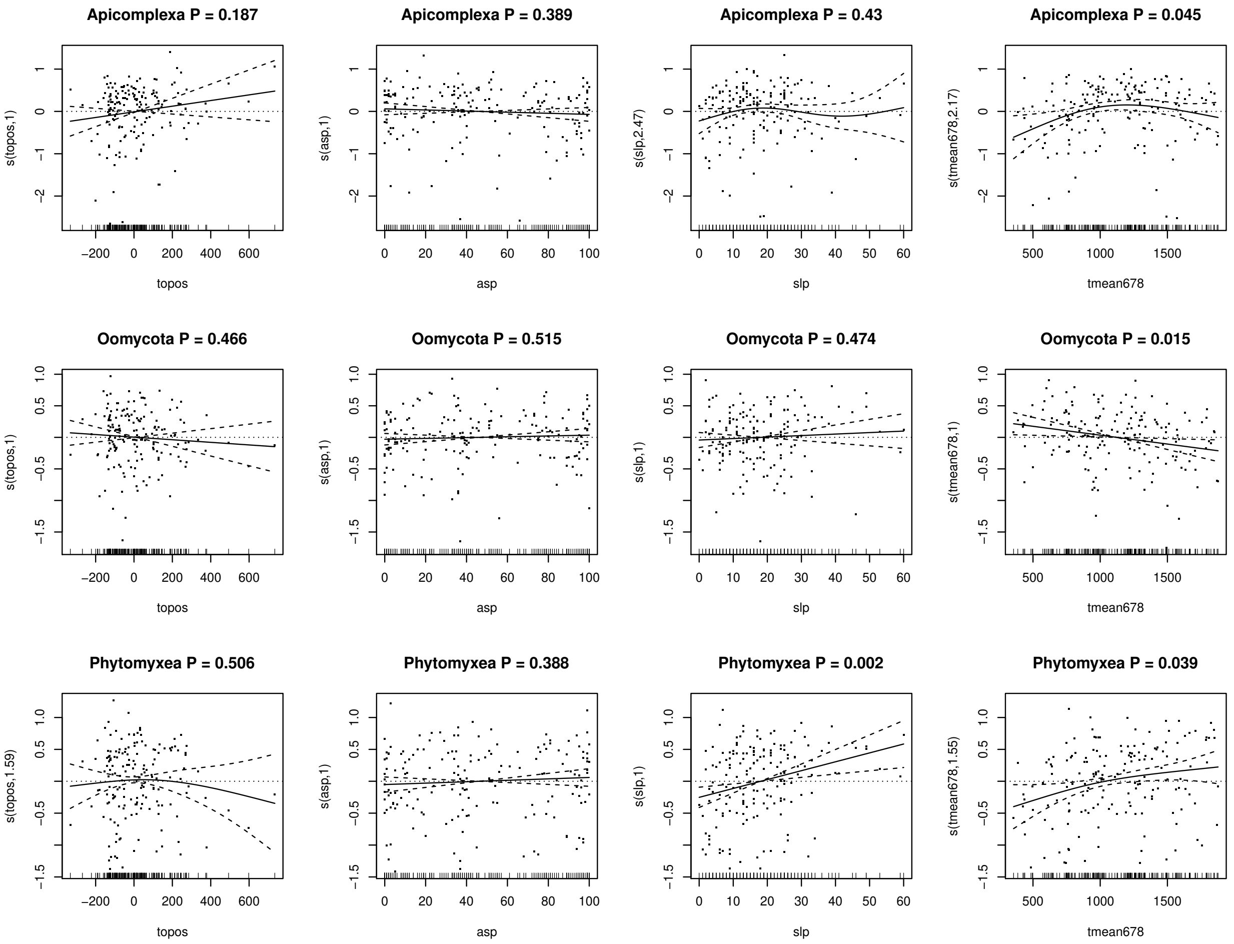

**Figure S4.2**: continuation

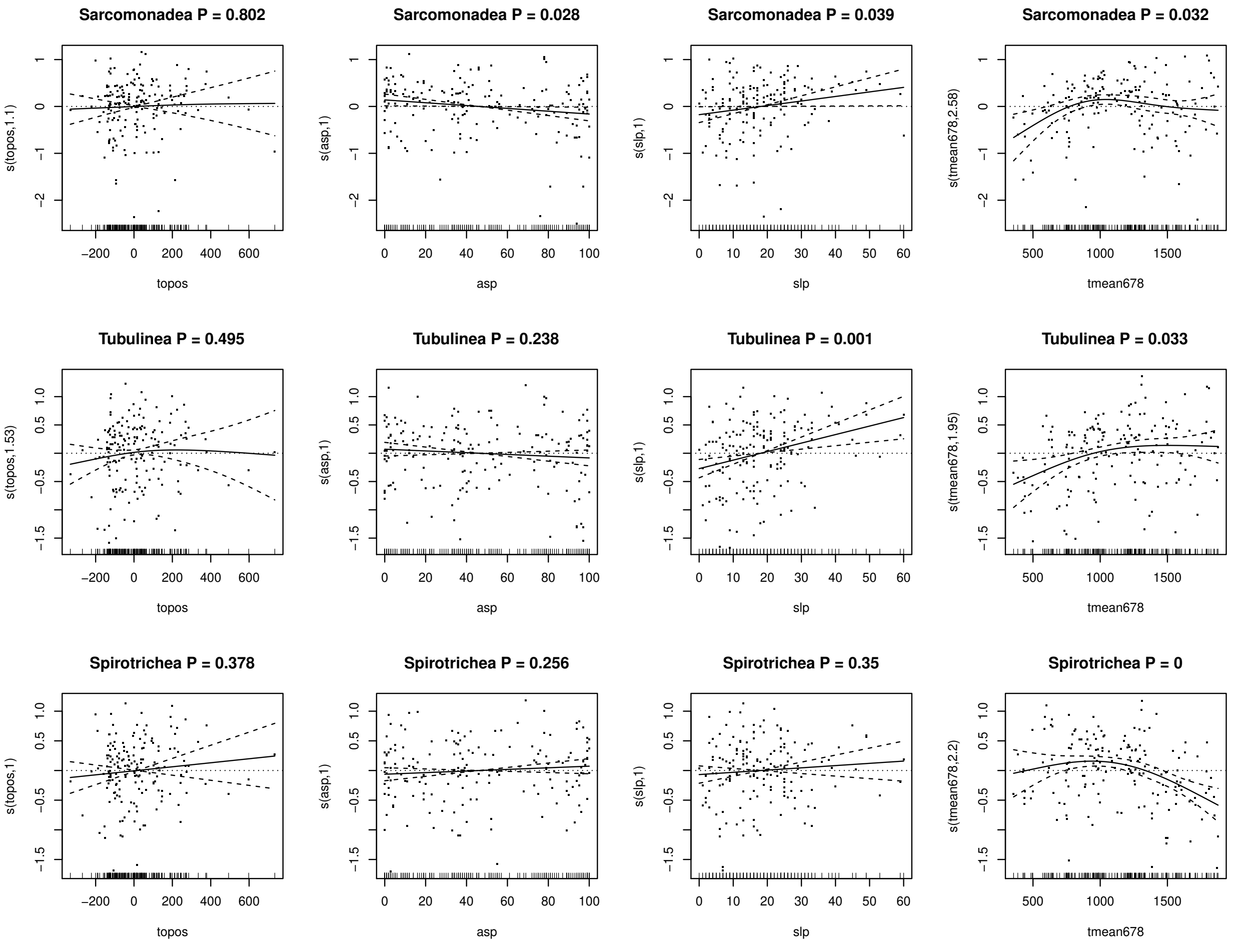

**Figure S4.2**: continuation

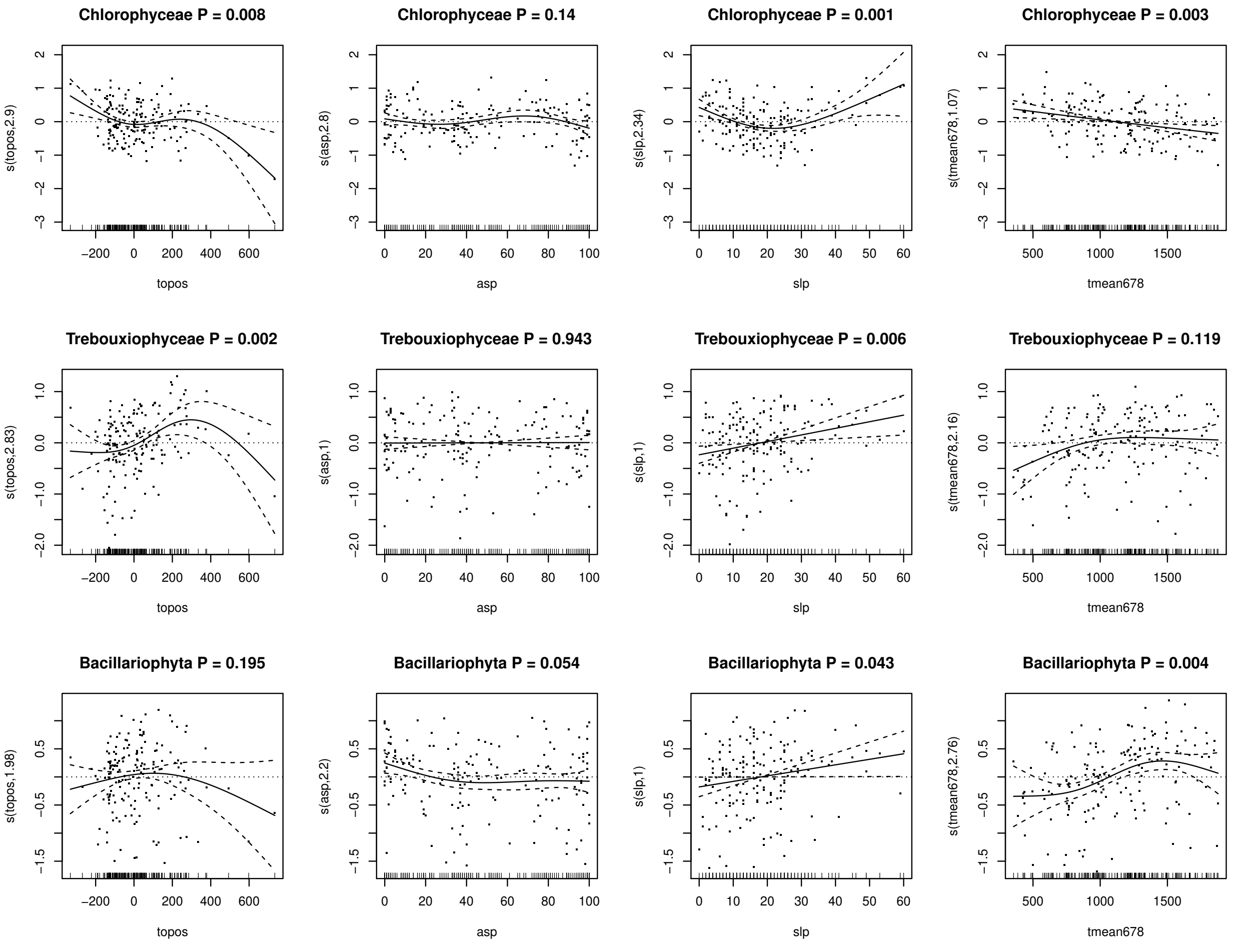
